# A reference genome for *Trichogramma kaykai*: A tiny desert-dwelling parasitoid wasp with competing sex-ratio distorters

**DOI:** 10.1101/2024.11.22.624848

**Authors:** Jack Culotta, Amelia RI Lindsey

**Affiliations:** Department of Entomology, University of Minnesota, Saint Paul, MN, USA 51108

**Keywords:** *Wolbachia*, sex ratio, selfish genetic element, symbiosis, B chromosome, *Trichogramma kaykai*

## Abstract

The tiny parasitoid wasp *Trichogramma kaykai* inhabits the Mojave Desert of the southwest United States. Populations of this tiny insect variably host up to two different sex-distorting genetic elements: (1) the endosymbiotic bacterium *Wolbachia* which induces the parthenogenetic reproduction of females, and (2) a B-chromosome, “Paternal Sex Ratio” (PSR), which converts would-be female offspring to PSR-transmitting males. We report here the genome of a *Wolbachia*-infected *Trichogramma kaykai* isofemale colony KSX58. Using Oxford Nanopore sequencing we produced a final genome assembly of 203 Mbp with 45x coverage, consisting of 213 contigs with an N50 of 1.9 Mbp. The assembly is quite complete, with 91.41% complete BUSCOs recovered: a very high score for Trichogrammatids that have been previously characterized for having high levels of core gene losses. We also report a complete mitochondrial genome for *T. kaykai,* and an assembly of the associated *Wolbachia*, strain *w*Tkk. We identified copies of the parthenogenesis-inducing genes *pifA* and *pifB* in a remnant prophage region of the *w*Tkk genome. The *Trichogramma kaykai* assembly is the highest quality genome assembly for the genus to-date and will serve as a great resource for understanding the evolution of sex and selfish genetic elements.

## INTRODUCTION

*Trichogramma* wasps (Hymenoptera: Trichogrammatidae) are some of the smallest animals on the planet (Polilov 2015). The genus contains more than 200 described species: all parasitoids that complete their development within the eggs of other insects (Burks et al. 2024; Pinto 2006). Trichogrammatid research has largely focused on (1) their application as biological control agents of insect pests (Knutson 1998; Cherif et al. 2021), (2) innovations associated with extreme miniaturization (Polilov 2012), and (3) sex allocation, especially due to relationships with sex-distorting elements (Stouthamer et al. 1990; Stouthamer and Kazmer 1994; Russell and Stouthamer 2010). The most common sex-ratio distorter is the intracellular, maternally transmitted bacterium *Wolbachia,* a common associate of many arthropods and nematodes (Kaur et al. 2021). In *Trichogramma*, most *Wolbachia* strains are “parthenogenesis-inducing” (PI), and enable the asexual reproduction of females (i.e., “thelytokous parthenogenesis”) (Stouthamer et al. 1990; Stouthamer et al. 1993; Ma and Schwander 2017).

To date all instances of microbe-mediated PI are in animals with haplodiploid sex determination (Ma and Schwander 2017; Verhulst et al. 2023). Under haplodiploidy (and without PI- *Wolbachia*) males typically develop from unfertilized (i.e., haploid) eggs, and females are typically derived from fertilized, diploid, eggs (De La Filia et al. 2015). PI-*Wolbachia* diplodize the unfertilized eggs, resulting in a female (Stouthamer and Kazmer 1994). In one species with PI-*Wolbachia*, *Trichogramma kaykai* (Figure 1A-B), a second sex-distorter is sometimes present: a supernumerary B-chromosome, “Paternal Sex Ratio” (PSR) (van Vugt et al. 2003; Stouthamer et al. 2001). PSR achieves the opposite outcome of *Wolbachia*’s PI: haploid males with PSR mate, and any fertilized eggs develop into more PSR-transmitting males (Van Vugt et al. 2009). PSR facilitates destruction of the paternal genome (except for itself), resulting in a haploid embryo (the maternal copy) and the untouched PSR chromosome. In populations where *Wolbachia* and PSR are present, a curious pattern of reproduction is present: males are derived from fertilized eggs (with PSR-containing sperm), and females are derived from unfertilized eggs (with PI-*Wolbachia*) (Figure 1C). Unlike many other PI-*Wolbachia* systems where PI is accompanied by a decay of sexual function (Stouthamer et al. 2010; Russell and Stouthamer 2011; Jeong and Stouthamer 2005; Stouthamer and Mak 2002; Gottlieb and Zchori-Fein 2001), *Trichogramma kaykai* are easily cured of their *Wolbachia* in the lab, and readily return to a fully functional sexual form (Hohmann and Luck 2000; Hohmann et al. 2001; Miura and Tagami 2004; Russell et al. 2016). The PSR chromosome ensures males and sexual reproduction are maintained.

**Figure 1.**
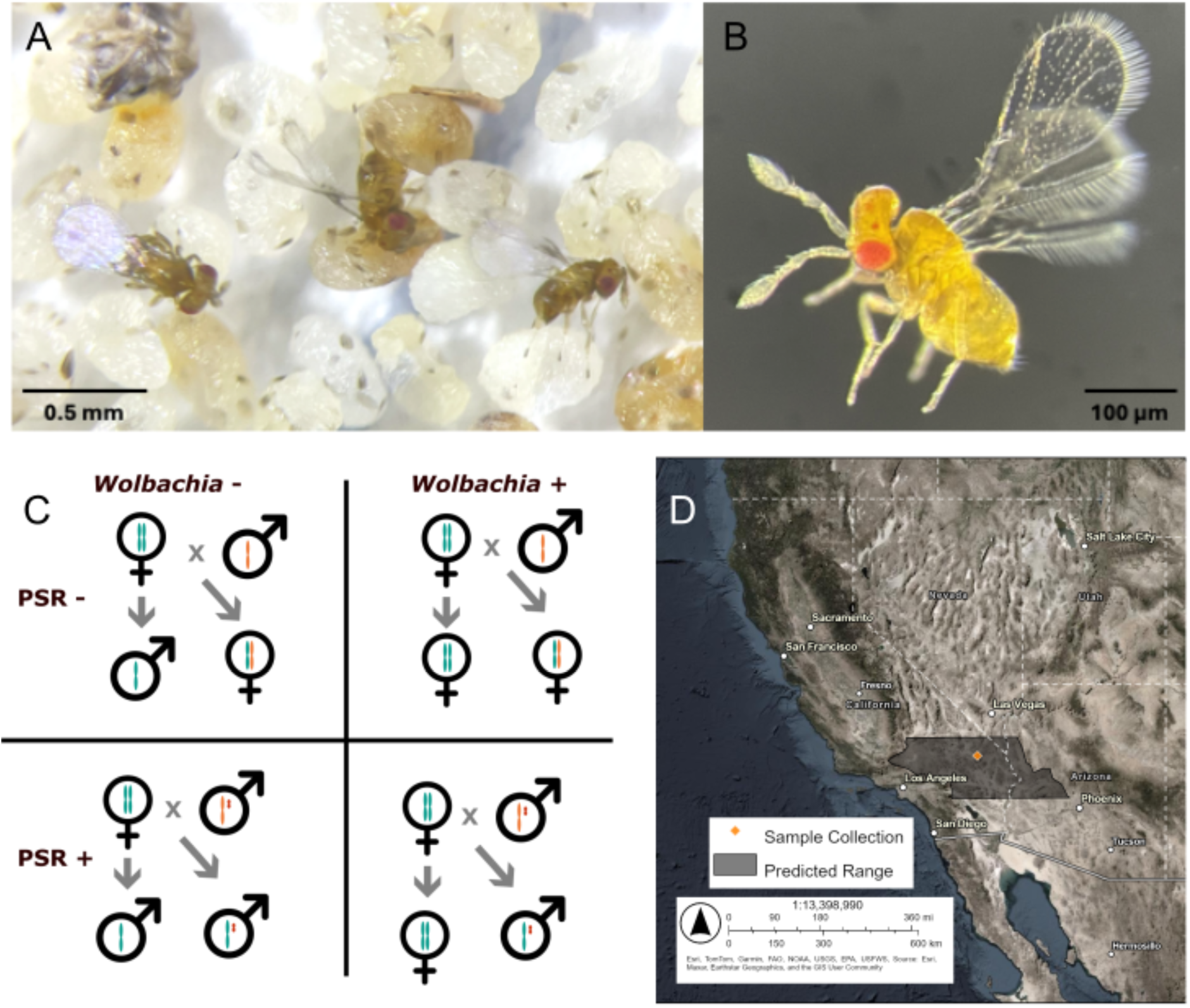
*Trichogramma kaykai* biology. **(A)** Three *Trichogramma kaykai* females ovipositing into host moth eggs (*Ephestia kuehniella*). **(B)** An exemplary specimen of *T. kaykai* (female). **(C)** Sex in *T. kaykai* is determined based on haplodiploidy, mediated by the presence or absence of *Wolbachia* (maternally transmitted) and the PSR chromosome (paternally transmitted). **(D)** The sample collection site for KSX58 and predicted geographic range of *Trichogramma kaykai*.

As host to PI-*Wolbachia* and PSR, *Trichogramma kaykai* is a valuable model for understanding the evolution of sex ratios and interactions between selfish genetic elements. This species was described in 1997 (Pinto et al.) and is native to the deserts of the Southwest United States (Figure 1D). We report a reference genome for an isofemale colony of *Trichogramma kaykai* from the Mojave Desert, plus the genome of its PI-*Wolbachia* strain, *w*Tkk. To our knowledge, there are currently no *Trichogramma kaykai* PSR chromosomes in culture, but this reference genome will aid in future efforts to understand how this selfish element alters chromosome dynamics and sex ratios.

## MATERIALS & METHODS

### Species Origin and Sampling Strategy

Genome sequencing and assembly was performed for *Trichogramma kaykai* line “KSX58”, an isofemale laboratory culture. A single unmated *Wolbachia*-infected, thelytokous female was reared out of a parasitized *Apodemia mormo* egg collected off an *Eriogonum inflatum* stem and used to initiate an isofemale line. The founding female was collected in May 2010 in Kelso, CA, USA, by R. Stouthamer and J. Russell (Figure 1D). The colony has since been maintained in 5 ml glass culture tubes stopped with cotton, and kept at 25°C with a 12:12 light:dark cycle. Wasps are hosted every 12 days on sterilized *Ephestia kuehniella* eggs adhered to cardstock alongside a streak of honey. *Wolbachia* infection status was confirmed by PCR with *Wolbachia* specific “Wspec" primers (Werren and Windsor 2000), and *Trichogramma* species was confirmed by molecular identification (Stouthamer et al. 1999), both as detailed previously (Lindsey and Stouthamer 2017). To collect wasps for DNA extraction, freshly emerged females were allowed to crawl up into a sterile tube attached to the colony culture vial. The pool of wasps was flash frozen in liquid nitrogen and stored at -80°C for further processing.

### Geographic Range Map

Locations of *Trichogramma kaykai* are centered around the Southern Mojave Desert (Pinto et al. 1997; Russell et al. 2018; Tulgetske and Stouthamer 2012; Russell et al. 2016; Van Vugt et al. 2009; van Vugt et al. 2003). The predicted northern and southern boundaries of this species’ range were estimated from these observations. As it is assumed *Trichogramma kaykai* is restricted to desert habitat, the eastern and western borders of range are indicated by the Southern Mojave Desert and Northern Sonoran Desert. The map was generated in ArcGIS Online (www.arcgis.com).

### Sequencing Methods and Sample Preparation

DNA was extracted from 25 mg of whole insect tissues using the MagAttract High Molecular Weight kit (Qiagen), following manufacturer’s instructions. The DNA was concentrated to 25 uL using Sergi Lab Supplies magnetic beads and went through the PacBio SRE kit to deplete fragments shorter than 10kb. The sample was barcoded and library prepped with the ONT SQK-NBD114.24 kit. The libraries were sequenced on a P2 Solo instrument using PromethION 10.4.1 flow cells. Every 24 hours the libraries were recovered and flowcells were flushed with nuclease (EXP-WSH004 kit) and reloaded.

### Nuclear Genome Assembly, Curation, and Quality Control

Samples were originally basecalled within Minknow using ‘super accuracy’ mode with 5mC_5hmC modified base calling. Reads were then re-basecalled with dorado v.0.7.2 using basecall model dna_r10.4.1_e8.2_400bps_sup\@v5.0.0. Reads at least 5kb in length were maintained, processed with ‘dorado correct’, and used for generating an assembly with Hifiasm v.0.19.9 and default parameters. The genome was manually curated, and cytoplasmic genomes were identified through tblastn results implemented in Blobtools v.1.1.1 (Challis et al. 2020). Assemblies were assessed with Compleasm v.0.2.6 (Huang and Li 2023) with the hymenoptera lineage flag (‘-l hymenoptera’).

### Genomic Methylation

Methylation and hydroxymethylation of genomic DNA at 5’ cytosines (5mC and 5hmC) in a cytosine-guanine dinucleotide (CpG) context was determined from the basecalling information stored in the unmapped modBAM files (Flack et al. 2024). These were aligned to the final assembly using Minimap v.2.17 (Li 2016), converted to bedMethyl format with Modkit v.0.4.1 (https://github.com/nanoporetech/modkit), and the 5mC and 5hmC percentages were calculated with an AWK script.

### Trichogramma Phylogeny

A whole-genome phylogeny was reconstructed with SANS v.2.4_10, which uses a pangenomic approach to calculate splits in a phylogenetic tree (Rempel and Wittler 2021). SANS parameters included ‘--filter strict’ with an output Newick tree file and 100 bootstrap replicates. Taxa included the available *Trichogramma* genomes (for *Trichogramma brassicae*, which is represented by two assemblies, only GCA_902806795.1 was used; Table 1), and an outgroup species from a closely related family (Cruaud et al. 2024), *Phymastichus coffea* (Hymenoptera: Eulophidae) GCF_024137745.1. Tree topology was configured in FigTree v.1.4.4 (https://github.com/rambaut/figtree/) and annotated in Inkscape (https://www.inkscape.org).

**Table 1.** Available *Trichogramma* genome assemblies.

| Species | Accession | Size (bp) | Contig/<br>Scaffold Count* | N50<br>(bp)* | *Wolbachia*<br>Genome | Citation |
| --- | --- | --- | --- | --- | --- | --- |
| *Trichogramma brassicae* | GCA_902806795.1 | 235,386,796 | 1,570 (C) | 556,663 | No | Ferguson et al. (2020) |
| *Trichogramma brassicae* | GCA_030522885.1 | 203,810,232 | 87,792 (S) | 18,131 | No | Guinet et al. (2023) |
| *Trichogramma dendrolimi* | GCA_034770305.1 | 215,209,100 | 316 (S) | 1,412,680 | No | Zhang et al. (2023) |
| *Trichogramma evanescens* | GCA_902732785.1 | 213,671,129 | 146,286 (S) | 38,173 | No | N/A |
| *Trichogramma pretiosum* | GCA_000599845.3 | 187,641,947 | 925 (S) | 1,825,723 | wTpre (Lindsey et al. 2016) | Lindsey et al. (2018a) |
| *Trichogramma kaykai* | Processing | 203,423,343 | 213 (C) | 1,898,390 | wTkk | This study |

\*If assembly is scaffolded, metrics reported are for scaffolds and an (S) is indicated in the count column. If there are only contigs, those metrics are reported and (C) is indicated in the count column.

### Repeat Assembly Techniques

We identified and masked repetitive sequences in each genome. First a custom *de novo* repeat library was crated with RepeatModeler v.2.0.5 (Flynn et al. 2020) with the -LTRStruct parameter included. Then this library was used to mask the genome with RepeatMasker v.4.1.1 (Tarailo-Graovac and Chen 2009) with the -s (sensitive mode) parameter included.

### Gene Finding Methods

To annotate the *T. kaykai* genome, a soft masked genome was used for gene model prediction with Galba v.1.0.11 (Brůna et al. 2023), using the RefSeq annotations for *Trichogramma brassicae* (GCA_902806795.1), *Trichogramma pretiosum* (GCA_000599845.3), *Nasonia vitripennis* (GCA_009193385.2), *Copidosoma floridanum* (GCF_000648655.2), *Phymastichus coffea* (GCF_024137745.1) and *Ceratosolen solmsi marchali* (GCF_000503995.2) as references. The Galba pipeline was executed using Singularity with parameters to output a gff3 file. Summary statistics for the resulting gff3 file were computed with GAG v.2.0.1 (Geib et al. 2018). Split genes (those encoded across the ends of two contigs) were manually re-assigned gene identifiers as per NCBI best practices.

### Synteny Analysis

We identified conserved regions and mapped synteny between the *T. kaykai* and *T. pretiosum* genomes (Table 1) using D-GENIES webtool (https://dgenies.toulouse.inra.fr/run) (Cabanettes and Klopp 2018) employing Minimap v.2.28 (Li 2016), the “many repeats” flag, and the “hide noise” option.

### Mitogenome

A single circular contig was identified as the mitochondrial genome based on GC content, size, and coverage. Mitogenome annotation was completed with MITOS2 v.2.1.9 (Bernt et al. 2013; Donath et al. 2019) and the circular mitogenome was started at Cox1 per convention with rearrangement in SnapGene v.7.2. MITOS2 parameters were the RefSeq63 Metazoa reference and the invertebrate mitochondrial translation code. Manual curation of the control region and inferences of gene structure were made based on comparisons to other *Trichogramma* mitochondrial genomes (Chen et al. 2018).

### *Wolbachia* Strain *w*Tkk Genome

Four contigs were identified as a *Wolbachia* genome based on cumulative size and Blobtools results. Genome completeness was analyzed against the rickettsiales_odb10 database with Compleasm v.0.2.6 (Huang and Li 2023). Prophage regions and mobile elements were identified with VirSorter2 v.2.2.4 (Guo et al. 2021) and mobileOG-db v.1.0.1 (Brown et al. 2022), implemented in proksee (Grant et al. 2023)(https://proksee.ca/) with default parameters. To identify putative parthenogenesis-inducing genes (*pifs*) (Fricke and Lindsey 2024), we leveraged annotation and orthology data generated by Prokka v.1.14.6 (Seemann 2014) and OrthoFinder v.2.5.4 (Emms and Kelly 2019), implemented in the *Wolbachia* Phylogeny Pipeline (WHOP; https://github.com/gerthmicha/WHOP). Phylogenetic analysis was performed based on the clustering results from WHOP/OrthoFinder results. Single-copy orthologs were aligned with MAFFT L-INS-i v.7.487 (Katoh and Standley 2013), recombining genes were eliminated with PhiPack v.1.1 (Bruen and Bruen 2005), and alignments were concatenated for phylogenetic reconstruction in IQtree v.2.2.3 (Nguyen et al. 2015), run with model optimization and 1000 ultrafast bootstrap replicates.

## RESULTS & DISCUSSION

### Sequencing and Assembly

We generated 10.5 billion base pairs of nanopore sequencing data: a total of 1,543,039 reads with a read N50 of 13,472 (Supplementary Table S1). A draft assembly from reads longer than 5,000 base pairs was generated with HiFiasm which produced a 204.6 Mbp assembly contained in 226 contigs. A combination of coverage, GC%, and blast hits from BlobTools results were used to identify non-nuclear contigs and curate the assembly. After removing spurious and contaminant contigs (Supplemental Table S2), and extracting the *Wolbachia w*Tkk and mitochondrial genomes, the final assembly was 203.4 Mbp in 213 contigs, with an average of 45x coverage (Table 2). The *Trichogramma kaykai* assembly falls in the middle of the size range for the genus, (187.6 Mbp in *T. pretiosum* to 235.4 Mbp in one of the *T. brassicae* (Table 1). Additionally, this size closely aligns with a flow cytometry-based estimate of 216 Mbp for a different colony of *T. kaykai*, “LC19-1” (van Vugt et al. 2005). The GC% of *Trichogramma* genomes appears to be highly conserved, with all at 40%.

**Table 2.**
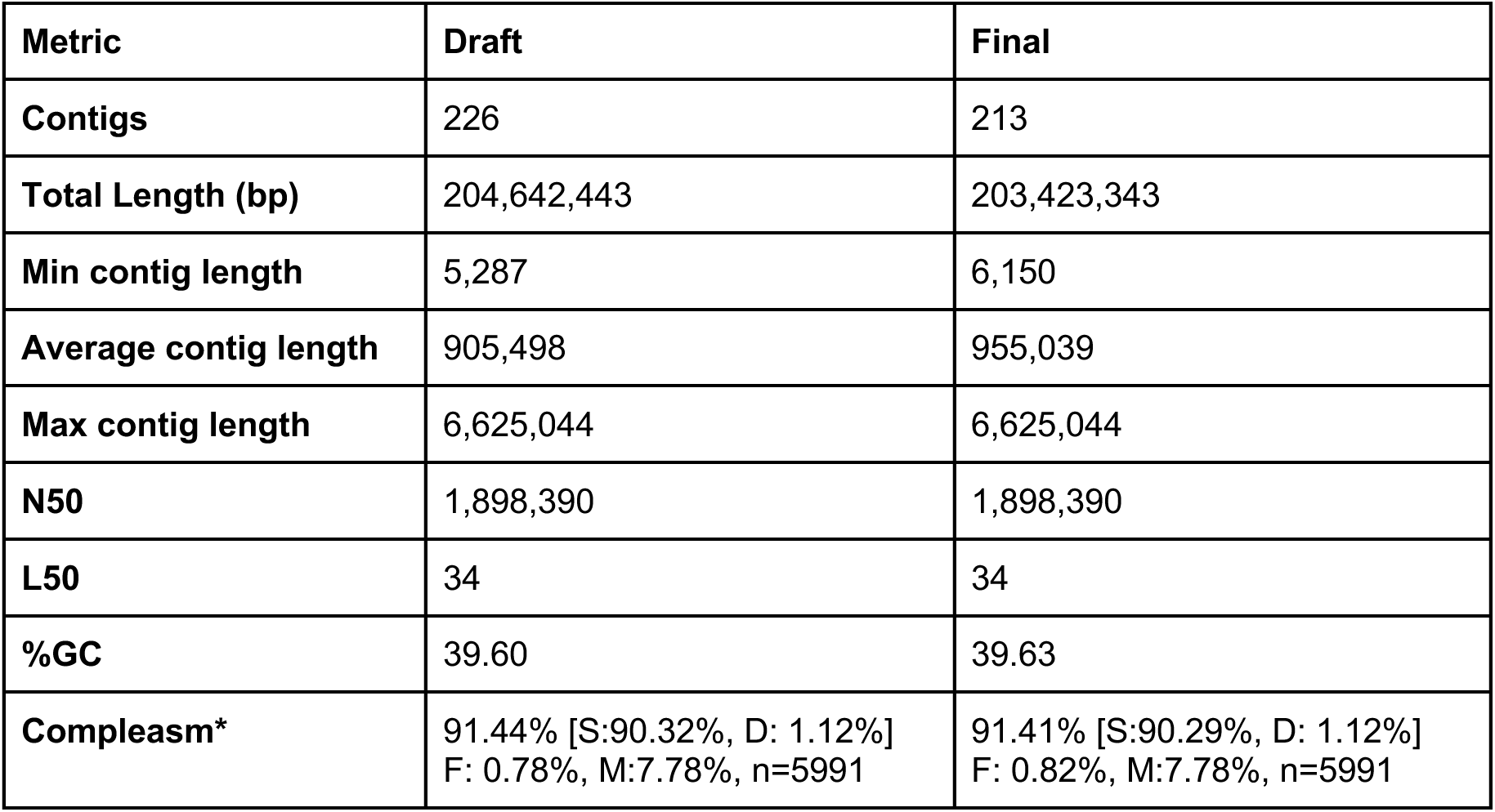
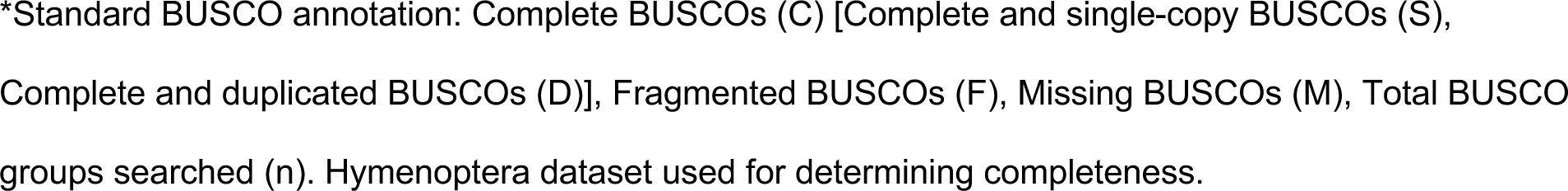
*Trichogramma kaykai* genome assembly statistics.

| Metric | Draft | Final |
| --- | --- | --- |
| Contigs | 226 | 213 |
| Total Length (bp) | 204,642,443 | 203,423,343 |
| Min contig length | 5,287 | 6,150 |
| Average contig length | 905,498 | 955,039 |
| Max contig length | 6,625,044 | 6,625,044 |
| N50 | 1,898,390 | 1,898,390 |
| L50 | 34 | 34 |
| %GC | 39.60 | 39.63 |
| Compleasm* | 91.44% [S:90.32%, D: 1.12%]<br>F: 0.78%, M:7.78%, n=5991 | 91.41% [S:90.29%, D: 1.12%]<br>F: 0.82%, M:7.78%, n=5991 |

\*Standard BUSCO annotation: Complete BUSCOs (C) [Complete and single-copy BUSCOs (S), Complete and duplicated BUSCOs (D)], Fragmented BUSCOs (F), Missing BUSCOs (M), Total BUSCO groups searched (n). Hymenoptera dataset used for determining completeness.

Quality assessments indicate that the genome assembly is quite complete, with 91.41% of Hymenopteran BUSCO loci present as complete coding sequences (Table 2). These metrics are on par with other well-assembled *Trichogramma* genomes (e.g., *T. pretiosum,* Hymenoptera BUSCO: C:92.7%[S:90.9%,D:1.8%],F:0.8%,M:6.5%,n:5991)(Lindsey et al. 2018a). Comparative genomics of *T. pretiosum* relative to other hymenopterans indicated that these wasps have undergone a large number of core gene losses and have highly accelerated rates of protein evolution (Lindsey et al. 2018a) so we do not expect BUSCO scores close to 100% even for a “perfect” assembly.

### Genome Methylation

We determined 5’ methylation at cytosines in a CpG context based on the direct sequencing basecalls. Less than 1% of CpGs were methylated: 0.67% of CpGs had 5mC (methyl) modifications and 0.18% had 5hmC (hydroxymethyl) modifications. While this is a low level of methylation as compared to vertebrates, this is not atypical for insects (Hunt et al. 2013). Importantly, this level of methylation closely mirrors the number of methylated CpG sites identified in *T. pretiosum* using bisulfite sequencing (Lindsey et al. 2018a; Wu et al. 2020).

### Analysis of Repetitive DNA

*Trichogramma kaykai* is sister to all other *Trichogramma* species with published genomes (Figure 2A). Across the genus, repetitive content appears to be relatively conserved. Repetitive sequences account for between 17.9% - 29.39% of the total genome lengths (Table 1, Figure 2B, Supplemental Table S3). This is in contrast to the outgroup species, *Phymastichus coffea* (Hymenoptera: Eulophidae), that has a 421 Mbp genome with more than half (57.21%) attributed to repetitive sequences (Figure 2B, Supplemental Table S3). Across *Trichogramma*, the majority of repetitive sequences are unclassified. In *T. kaykai*, 4% of the genome is derived from retroelements, <1% from DNA transposons, and around 3% of the genome is simple and low complexity repeats (Table 3). We then assessed the level of synteny between *T. kaykai* and *T. pretiosum* by cross-mapping similar genomic sequences with D-GENIES (Figure 3). A large proportion (60.34%) of the *T. kayaki* genome shares 50-75% identity with *T. pretiosum*, and there are high levels of synteny across the two assemblies (Figure 3).

**Figure 2.**
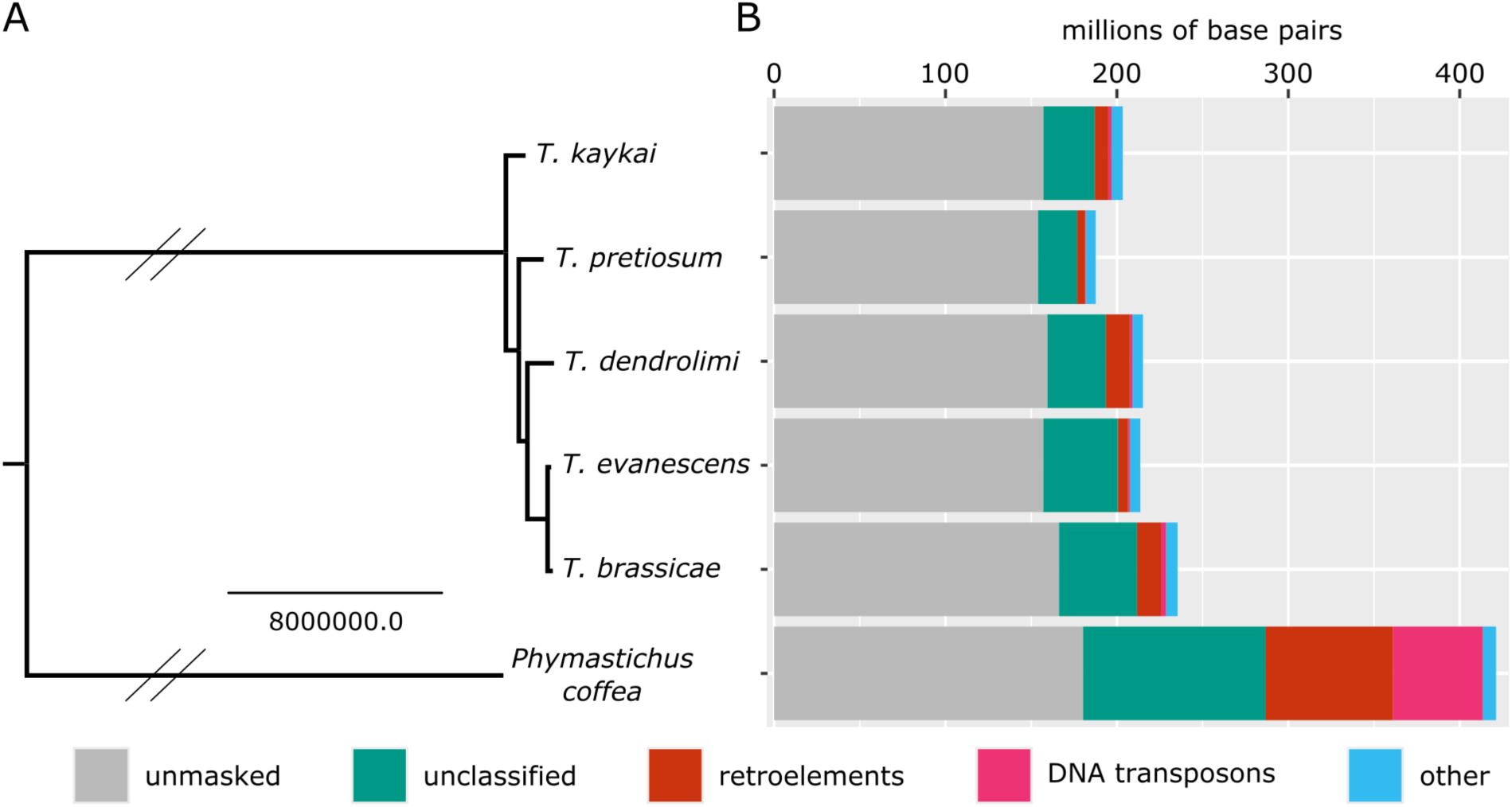
Comparative genomics of *Trichogramma.* **(A)** Whole genome phylogeny of five *Trichogramma* species and outgroup *Phymastichus coffea* (Hymenoptera: Eulophidae). Double slashes indicate branches that were shortened to half their length for ease of visualization. **(B)** Repetitive content of each genome, corresponding to the taxa in (A). “Other” includes rolling circles, simple repeats, and low complexity repeats.

**Figure 3.**
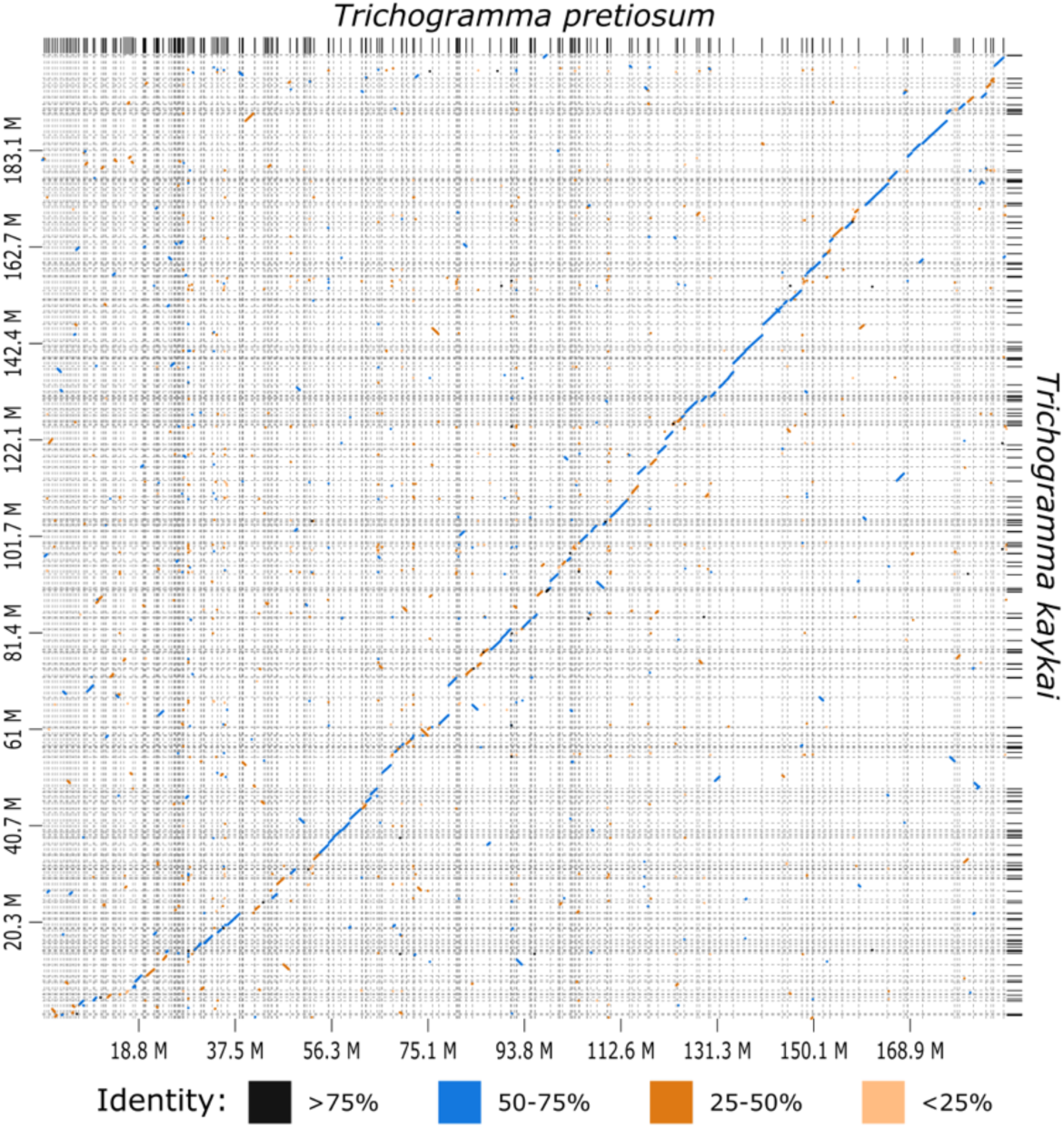
Synteny is highly conserved between *T. kaykai* and *T. pretiosum*. Dot plot indicating syntenic regions between *Trichogramma* genomes. Dots are colored according to percent identity.

**Table 3.** Interspersed repeats in *Trichogramma kaykai*.

| **Name** | **Number** | **Length (bp)** | **Percent (%)** |
| --- | --- | --- | --- |
| **Retroelements** | 8,591 | 8,132,788 | 4.00% |
| Penelope class | 203 | 76,919 | 0.04 % |
| LINE class | 5,232 | 5,084,665 | 2.50 % |
| L2/CR1/Rex | 674 | 423,279 | 0.21 % |
| R1/LOA/Jockey | 1,929 | 1,977,765 | 0.97 % |
| R2/R4/NeSL | 2,056 | 2,301,966 | 1.13 % |
| LTR class | 3,359 | 3,048,123 | 1.50 % |
| BEL/Pao | 484 | 436,375 | 0.21 % |
| Ty1/Copia | 497 | 368,036 | 0.18 % |
| Gypsy/DIRS1 | 2,378 | 2,243,712 | 1.10 % |
| **DNA transposons** | 2,812 | 1,863,150 | 0.92 % |
| hobo-Activator | 497 | 176,867 | 0.09 % |
| Tc1-IS630-Pogo | 370 | 118,645 | 0.06 % |
| **Rolling-circles** | 797 | 350,441 | 0.17 % |
| **Unclassified** | 110,502 | 29,820,645 | 14.66 % |
| **Total interspersed repeats** |  | 39,816,583 | 19.57 % |
| **Simple repeats** | 147,687 | 5,403,933 | 2.66 % |
| **Low complexity** | 15,687 | 704,136 | 0.35 % |
| **Bases masked** |  | 46,275,093 | 22.75 % |

### Genome Annotation

We annotated the *T. kaykai* genome using a set of protein sequences from other wasps in the superfamily Chalcidoidea as a reference and identified 20,798 genes (Table 4). These genes corresponded to 24,714 transcripts, with a mean of four exons per mRNA (Table 4). Compared to other *Trichogramma* species, this is a larger number of annotated genes (e.g., 13,395 in *T. pretiosum*, 16,905 in *T. brassicae*). However, this could be due to differences in annotation pipelines, and or, lineage-specific differences in the patterns of gene gain and loss.

**Table 4.**
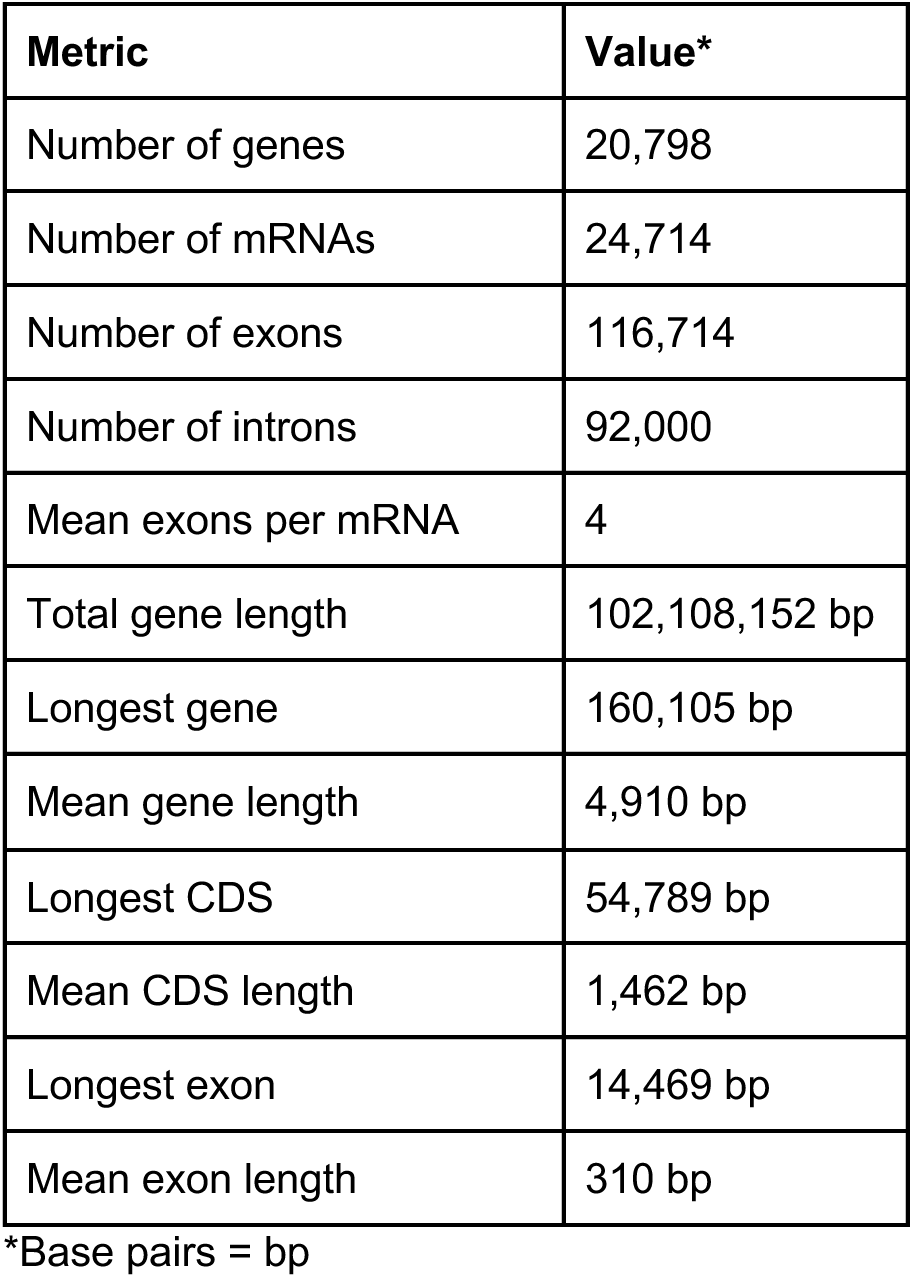
Annotation metrics for the *Trichogramma kaykai* genome.

### Mitogenome

We identified the mitogenome based on GC content (14.81%), size (16,399 bp), and coverage (3708x). Annotation revealed all expected mitochondrial tRNAs and coding genes (Figure 4). MITOS2 annotated a single large rRNA of only 712 bp and three regions (387, 49, and 38 bp) as small rRNAs. Comparison to other Trichogrammatid mitochondrial genomes indicated that the large rRNA annotation had been truncated on the 5’ end, and the small rRNA annotation had been fragmented (Figure 4), which is likely due to the extreme divergence of these mitochondrial sequences. A 878 bp region between the tRNAs for tryptophan (W) and methionine (M) corresponds to the putative control region identified in other Trichogrammatid mitochondrial genomes (Chen et al. 2018).

**Figure 4.**
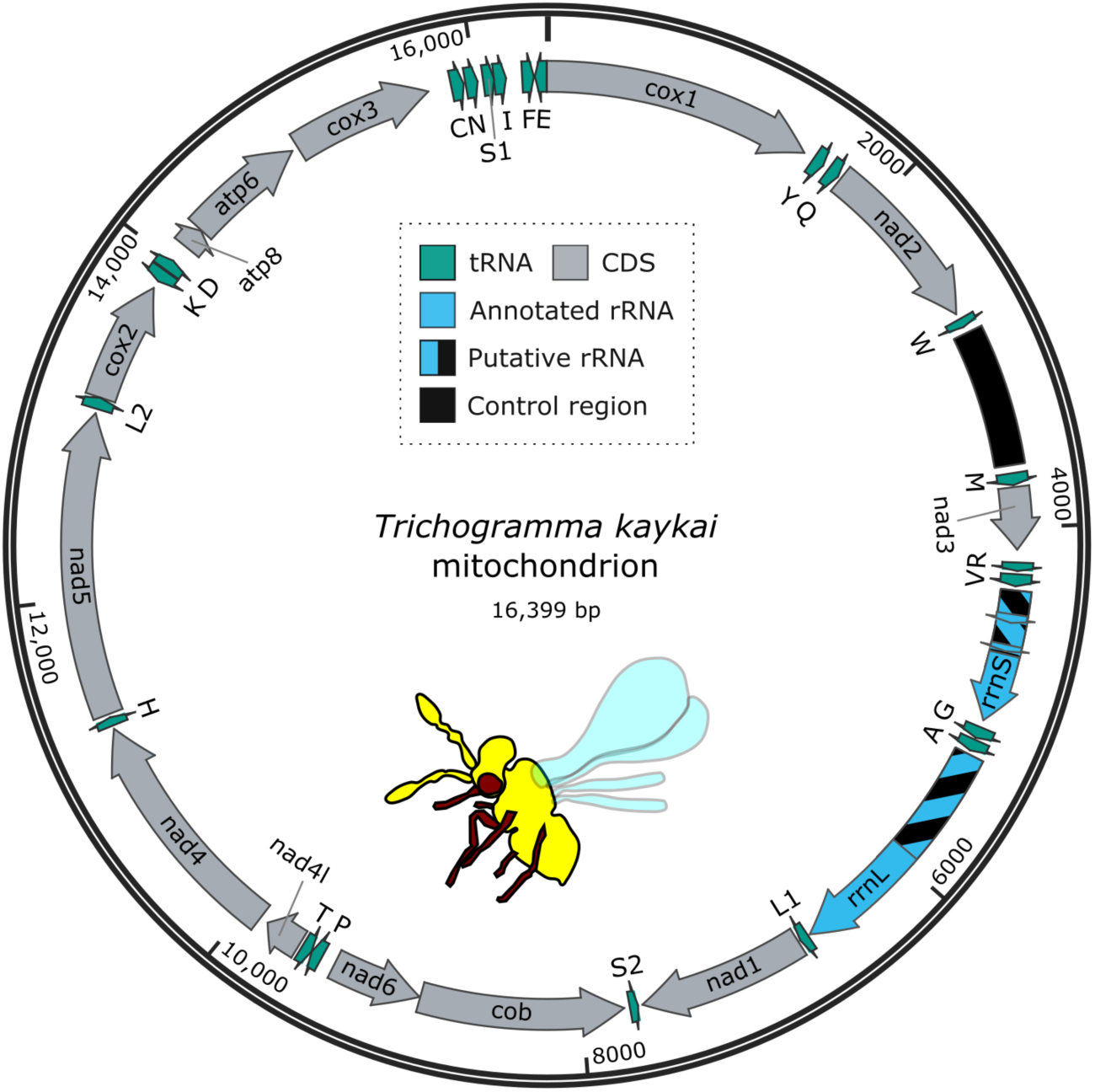
Mitochondrial genome of *Trichogramma kaykai*. Genes were annotated with MITOS2 (Bernt et al. 2013). Putative regions of rRNAs that were not correctly annotated by MITOS2 are indicated with stripes. The control region and the putative full length rRNAs were identified based on homology and gene order of other *Trichogramma* mitochondria (Chen et al. 2018). Transfer RNAs (tRNAs) are denoted by IPUC-IUB amino acid codes.

### Parthenogenesis-Inducing *Wolbachia* Strain *w*Tkk

We assembled a near-complete *Wolbachia* genome of the *w*Tkk strain: ∼1.12Mbp contained in four contigs, sequenced at 55X coverage (Table 5). Phylogenetic reconstruction revealed that *w*Tkk is in the “Supergroup B” clade of *Wolbachia*, and is sister to *w*Tpre, which infects *Trichogramma pretiosum* (Lindsey et al. 2016)(Figure 5A). The *w*Tkk and *w*Tpre genomes are similar in size: the *w*Tpre assembly (a single scaffold) is just slightly larger at 1,133,709 bp (Lindsey et al. 2016). We queried the *w*Tkk proteins to identify the recently identified parthenogenesis inducing factors, *pifA* and *pifB* (Fricke and Lindsey 2024). We identified a single copy of each gene in the *w*Tkk genome, encoded next to each other within a remnant prophage region (Figure 5B), as is typical of many other *Wolbachia* loci that induce host reproductive manipulations (Fricke and Lindsey 2024; LePage et al. 2017; Lindsey et al. 2018b; Shropshire et al. 2018; Perlmutter et al. 2019; Bordenstein and Bordenstein 2016). The *w*Tkk PifA protein was 67% identical to the PifA from *w*Tpre, and 30% identical to the PifA from *w*Lcla (another PI-*Wolbachia* infecting the parasitoid wasp *Leptopilina clavipes* [Hymenoptera: Figitidae])(Pannebakker et al. 2004). In contrast, PifB proteins were more conserved: *w*Tkk and *w*Tpre PifB were 93% identical, and *w*Tkk and *w*Lcla PifB shared 56% amino acid identity.

**Figure 5.**
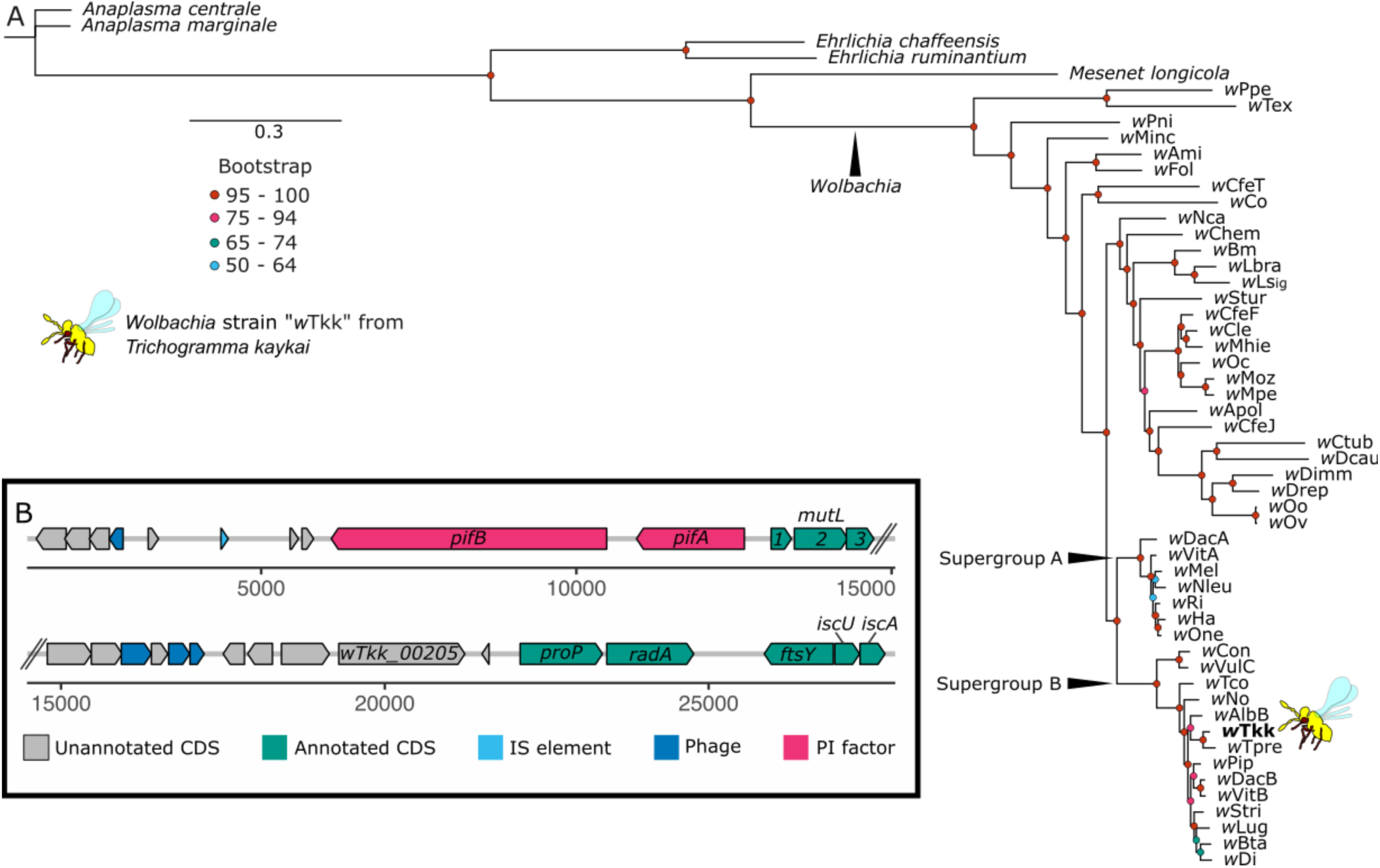
Parthenogenesis-inducing *Wolbachia* strain *w*Tkk. **(A)** Maximum likelihood-based phylogeny of *Wolbachia* strains and Rickettsiales outgroups based on 78 core, single-copy, protein coding genes (a total of 30,477 aligned amino acid sites). **(B)** Gene models for a predicted remnant prophage region that contains the parthenogenesis factors *pifA* and *pifB.* Three tandem CDS were annotated as *mutL*, which is likely a pseudogenization of *mutL* due to nonsense mutations and fragmentation of the coding region into multiple open reading frames. Abbreviations: insertion element (IS), parthenogenesis inducing factor (*pif*), coding sequence (CDS).

**Table 5.**
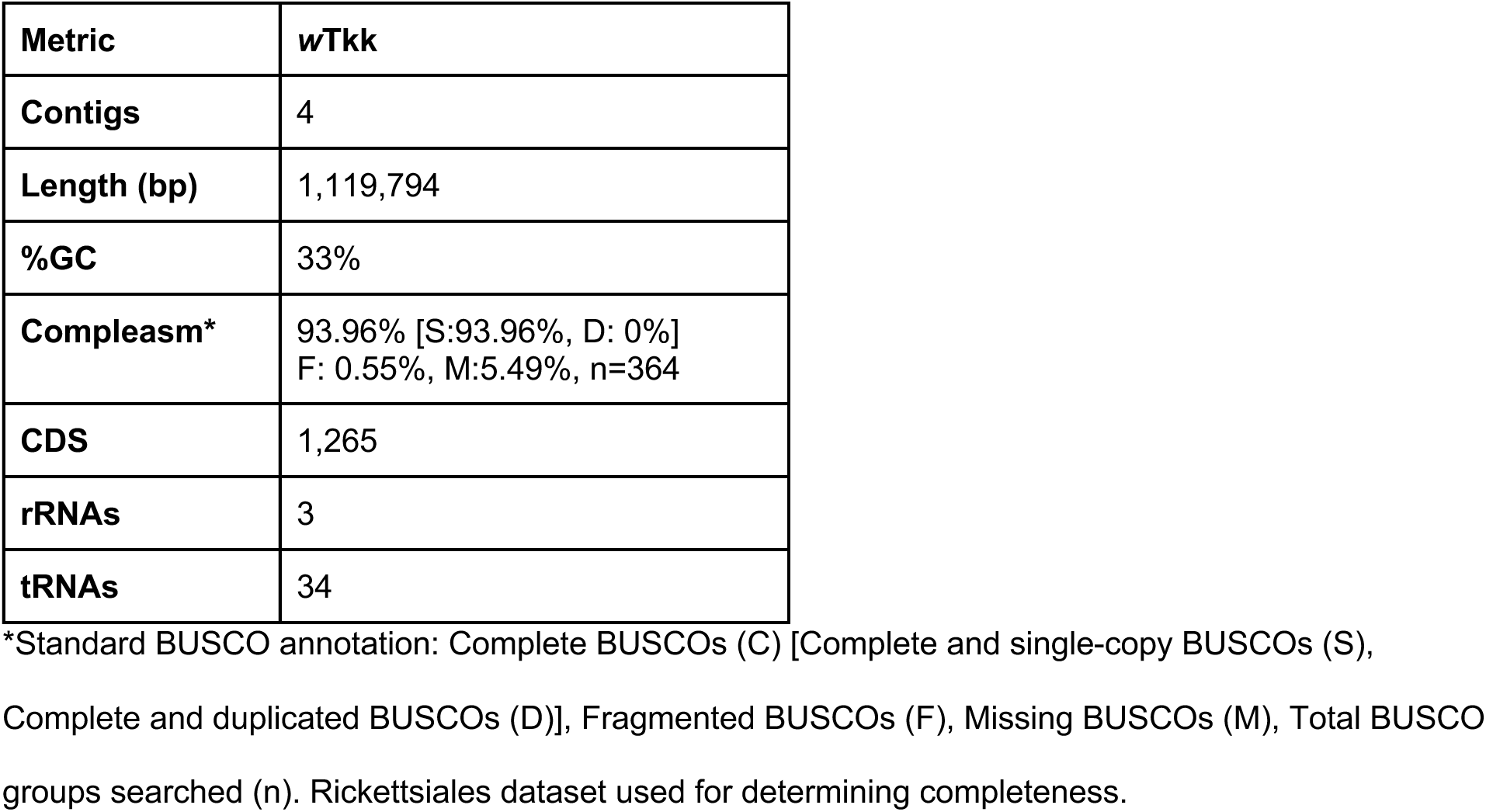
Wolbachia strain wTkk genome assembly and annotation.

| Metric | wTkk |
| --- | --- |
| Contigs | 4 |
| Length (bp) | 1,119,794 |
| %GC | 33% |
| Compleasm* | 93.96% [S:93.96%, D: 0%]<br>F: 0.55%, M:5.49%, n=364 |
| CDS | 1,265 |
| rRNAs | 3 |
| tRNAs | 34 |

\*Standard BUSCO annotation: Complete BUSCOs (C) [Complete and single-copy BUSCOs (S), Complete and duplicated BUSCOs (D)], Fragmented BUSCOs (F), Missing BUSCOs (M), Total BUSCO groups searched (n). Rickettsiales dataset used for determining completeness.

## SUMMARY

We report here a high-quality assembly for the parasitoid wasp *Trichogramma kaykai* along with genomes for its mitochondrion and associated *Wolbachia* strain, *w*Tkk. There are five other *Trichogramma* genomes currently available on NBCI: one each from the *Trichogramma* species *pretiosum, dendrolimi,* and *evanescens*, and two assemblies for *Trichogramma brassicae*. These species are some of the more commonly available *Trichogramma* sold as biological control agents of lepidopteran pests (Knutson 1998; Cherif et al. 2021). The *Trichogramma kaykai* assembly reported here is arguably the highest quality assembly available for the genus, as it has the lowest number of contigs and the highest N50 (Table 1). To date, all *Trichogramma* species assayed for karyotype have a haploid genome of five chromosomes (2n=10) (Gokhman and Quicke 1995; Van Vugt et al. 2009; Gokhman 2020; Gokhman et al. 2017; Farsi et al. 2020). While chromosome number and approximate genome size are conserved, there do appear to be species-specific differences in chromosome morphometrics (e.g., centromere location, arm lengths, chromosome sizes) (Farsi et al. 2020; Gokhman et al. 2017; Gokhman 2020).

Of the *Trichogramma* genome sequencing efforts, one other reports a *Wolbachia* genome: strain *w*Tpre, from *T. pretiosum* (Lindsey et al. 2016). The two *Trichogramma*-infecting strains*, w*Tpre and *w*Tkk, are closely related members of the “Supergroup B” clade which contains a suite of other arthropod-infecting strains, including other parthenogenesis-inducers from a range of host insects (Scholz et al. 2020; Lindsey et al. 2016). While *Wolbachia* are maternally transmitted, across longer evolutionary time scales there is a significant amount of horizontal transfer, and often sister strains infect distantly related hosts (Bailly-Bechet et al. 2017; Scholz et al. 2020). However, the PI-*Wolbachia* infecting *Trichogramma* appear to have a single origin (Poorjavad et al. 2012; Schilthuizen and Stouthamer 1997; Almeida and Stouthamer 2017). These PI-*Wolbachia* still undergo host switching (i.e., there is no co-cladogenesis) (Huigens et al. 2000; Huigens et al. 2004; Almeida and Stouthamer 2017), but that this clade of *Wolbachia* seem restricted to a single host genus makes them an interesting case study for host adaptation and the evolution of their PI effector proteins.

In addition to the PI-*Wolbachia* present in *T. kaykai*, the PSR chromosome found in some males offers another opportunity to understand the evolution of sex ratio distortion (Zhang and Ferree 2024). One other such PSR chromosome has been described: in the parasitoid wasp *Nasonia vitripennis* (Hymenoptera: Pteromalidae)(Werren 1991; Nur et al. 1988). The PSR chromosomes from these two wasp species appear to have independent origins, albeit a very similar paternal genome elimination phenotype (van Vugt et al. 2003; Zhang and Ferree 2024). Curiously, both PSR chromosomes seem to have originated from hybridization events in which chromosomal regions with abundant repetitive elements were transferred in via a close relative (McAllister and Werren 1997; van Vugt et al. 2005; Van Vugt et al. 2009). In contrast to *T. kaykai*, *Nasonia* are not known to host any PI symbionts (Beukeboom and Van De Zande 2010). However, some *Nasonia vitripennis* do host male-killing bacteria: *Arsenophonus nasoniae* (Gherna et al. 1991; Ferree et al. 2008). The PSR chromosomes are likely playing a key role in male-rescue which balances the male-eliminating cytoplasmic factors in both systems (either elimination by conversion to female via PI-*Wolbachia* in *Trichogramma*, or, elimination via death via *Arsenophonus* in *Nasonia*). The *Wolbachia*-infected line of *T. kaykai* reported here will enable the long-term maintenance of PSR chromosomes in the lab, and in the future, we hope to re-collect PSR-containing males from the native range to better understand the evolution of these selfish genetic elements.

## Data Availability

This Whole Genome Shotgun project has been deposited at DDBJ/ENA/GenBank under the BioProject accession PRJNA1150630. BioSample accessions for *Trichogramma kaykai* and *Wolbachia* strain *w*Tkk are SAMN43292057 and SAMN43292058, respectively. Sequencing reads are deposited under SRR30339640. Genome assemblies and annotations for the *Trichogramma kaykai* nuclear genome, *Trichogramma kaykai* mitochondrial genome, and *Wolbachia* strain *w*Tkk are currently processing. Supplemental materials are available on the GSA Figshare portal and include: (A) Table S1. Nanopore sequencing statistics, (B) Table S2. Details on draft assembly curation, (C) Table S3. Comparison of interspersed repeats between *Trichogramma kaykai* and related species, (D) File S1. *Trichogramma kaykai g*enome annotations in GFF3 format, and (E) File S2. Analysis notebook with bioinformatics workflows and scripts. A voucher of the *Trichogramma kaykai* KSX58 colony is available at the University of California Riverside Insect Collection: UCRC_ENT00496298.

## Acknowledgements

We thank Chris Faulk and Carrie Walls of Decorative Genomics (https://decogenomics.com/) for providing sequencing services. Many thanks to Richard Stouthamer for gifting ARIL the KSX58 colony. We acknowledge that the initial collection of insects to start a colony occurred on the traditional land of the Mojave People, and we are deeply grateful to the people who have stewarded the land throughout generations.

## Author Contributions

ARIL provided samples, funding, analytical guidance, and performed some analyses. JC performed molecular work and bioinformatic analyses. JC and ARIL co-wrote the manuscript and created figures. Both authors read and approved the manuscript prior to submission.

